# Effective reserve: a latent variable to improve outcome prediction in stroke

**DOI:** 10.1101/192823

**Authors:** Markus D. Schirmer, Mark R. Etherton, Adrian V. Dalca, Anne-Katrin Giese, Lisa Cloonan, Ona Wu, Polina Golland, Natalia S. Rost

## Abstract

**Background:** Prediction of functional outcome after stroke based on initial presentation remains an open challenge, suggesting that an important aspect is missing from these prediction models. There exists the notion of a protective mechanism called brain reserve, which may be utilized to understand variations in disease outcome. In this work we expand the concept of brain reserve (effective reserve) to improve prediction models of functional outcome after acute ischemic stroke (AIS).

**Methods:** Consecutive AIS patients with acute brain MRI (<48 hours) were eligible for this study. White matter hyperintensity and acute infarct volume were determined on T2 fluid attenuated inversion recovery and diffusion weighted images, respectively. Modified Rankin Scale scores (mRS) were obtained at 90 days post-stroke. Effective reserve (eR) was defined as a latent variable using structural equation modeling by including age, systolic blood pressure, and intracranial volume measurements.

**Results:** Of 453 AIS patients (mean age 66.6±14.7 years), 36% were male and 311 hypertensive. There was inverse association between eR and 90-day mRS (path coefficient -0.18±0.01, p<0.01). As compared to a model without eR, correlation between predicted mRS and observed mRS improved in the eR-based model (Spearman’s ρ 0.29±0.18 versus 0.15±0.17, p<0.001). Furthermore, hypertensive patients exhibited lower eR (p<10^−6^).

**Conclusion:** Using eR in prediction models of stroke outcome is feasible and leads to better model performance. Furthermore, higher eR is associated with more favorable functional post-stoke outcome and might correspond to an overall better vascular health.

## Introduction

Early prediction of stroke risk and a better understanding of the underlying disease processes may lead to effective prevention strategies for adverse cognitive and functional stroke outcomes, thereby improving patients’ quality of life and reducing the economic burden on families and society [1]. However, mechanisms of post-stroke recovery are complex and not well understood.

Among these, the notion of biological protective mechanisms intrinsic to the brain’s ability to withstand insults has been linked to better outcomes after disease onset [2–4]. This innate resilience to insults is in part supported by the observation that some subjects experience a better outcome than expected given their clinical presentation, and is increasingly referred to as “brain reserve” (BR) [5,6]. BR has been proposed as an indirect measure of brain health [7] and linked to brain volume [8]; however, understanding of the biological underpinnings of such a mechanism remains limited.

In stroke, a theoretical construct such as BR could improve prediction models of post-stroke outcomes; however, theories describing such mechanism are lacking. Prior non-stroke studies that modeled BR as a latent variable in structural equation modeling (SEM) analysis have, for example, demonstrated that BR is positively correlated with white matter hyperintensity volume (WMHv) in healthy elderly, while controlling for speed/executive or language function [9]. This suggests that patients with higher BR can withstand a greater burden of cerebral microvascular disease (larger WMHv) before significant deterioration in cognitive function occurs.

SEM is commonly used to confirm whether a model is supported by the data and can be seen as a combination of multiple regression and confirmatory factor analysis, in which latent variables are introduced to model properties of the system that are not measured or cannot be observed directly[10]. Given that WMHv is a validated risk factor for poor outcomes after stroke [11], we hypothesize that a model which accounts for an intrinsic (unmeasured) biological protective mechanism (such as BR), while accounting for known risk factors such as WMHv, might improve prediction of stroke outcome [12,13].

Furthermore, in this work we expand on the idea of BR, by defining a latent variable we call “effective reserve” (eR), which models the resilience of the brain after negative effects, e.g. of the brain injury, have been taken into account. We hypothesize that accounting for the brain’s intrinsic ability to withstand the burden of chronic (pre-stroke) and acute (acute ischemia) insults will improve our ability to predict functional outcomes after stroke. To test this hypothesis, we examine: (a) whether a model that includes eR can describe functional stroke outcome as captured by the modified Rankin scale score (mRS) at 90 days post-stroke[14] better than a linear model based on the observed variables only; and (b) the biological underpinnings of what eR may represent in relation to overall vascular health of the patient based on pre-stroke diagnosis of hypertension.

## Methods

### Standard protocol approvals, registrations, and patient consent

At the time of enrollment in this study, informed written consent was obtained from all participating subjects or their surrogates. The use of human subjects in this study was approved by the Institutional Review Board.

### Study design, setting, and patient population

Patients presenting to the emergency department at our hospital between 2003 and 2011, who were over 18 years of age and showed signs and symptoms of acute ischemic stroke (AIS), were eligible for enrollment in the institutional ischemic stroke cohort. In this analysis, we included subjects with (a) acute cerebral infarct lesions confirmed by diffusion weighted imaging (DWI) scans obtained within 48 hours of symptom onset and (b) T2 fluid-attenuated inversion recovery (T2-FLAIR) sequences available for WMHv analysis (total n=481 subjects).

### Clinical variables and outcome assessment

All clinical variables including demographics, past medical history, and vital signs were obtained on admission. Patients and/or surrogates were interviewed directly and medical records were reviewed, as necessary.

All variables were assessed for outliers. Patients with systolic blood pressure (SBP) below 50 (20 patients) or above 250 (4 patients), as well as intracranial volume (ICV) below 12 cm^3^ (4 patients) were excluded, resulting in a total of 453 subjects. Hypertensive patients were defined as those with past medical history of hypertension or those who were on anti-hypertensive medication (311/453, 68.7%). Hypertension status was missing for 72 of the patients.

Patients and/or their caregivers were interviewed in person or by telephone at 3-6 months after AIS to assess functional outcome (mRS). Recurrent cerebrovascular events, newly diagnosed medical conditions, and medication use were specifically assessed in this interview. If the patient could not be contacted, mRS was determined from chart review.

### Neuroimaging analysis

The standard AIS protocol included DWI (single-shot echo-planar imaging; one to five B0 volumes, 6 to 30 diffusion directions with b=1000 s/mm^2^, 1-3 averaged volumes) and T2 FLAIR imaging (TR 5000ms, minimum TE of 62 to 116ms, TI 2200ms, FOV 220-240mm). DWI data sets were assessed and corrected for motion and eddy current distortions [15]. WMHv was determined on the T2-FLAIR images using a semi-automated approach [16] in the hemisphere contralateral to the acute ischemic lesion (i.e., contralesional) and doubled to estimate the total WMH burden. Additionally, acute infarct volume was manually assessed on DWI (DWIv). ICV was calculated on T1 sagittal sequences using a previously validated method [17].

### Statistical analysis of clinical variables

Group differences in clinical variables between hypertensive and non-hypertensive patients are assessed using Wilcox and Fisher exact test for continuous and categorical variables, respectively. Inter-measure correlations are assessed using Spearman’s rank correlation coefficient. P-values of correlations are approximated based on t-/F-distributions, as described previously[18] and implemented in the R package Hmisc[19].

### Effective Reserve

In our model, eR is represented as a latent variable. We assume ICV to have a direct (positive) association with eR [9,20]. Additionally, age is believed to limit the brain’s capacity to counteract insults [21]. This reduced capacity to withstand the impact of an acute insult such as AIS might be reflected in long-term exposure to abnormally elevated blood pressure, as reflected in admission SBP. Therefore, we include age, SBP and ICV in our eR model, where eR is given by:

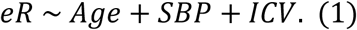

Moreover, we incorporate known links between age and WMHv [22–24], SBP and WMHv [22–25], WMHv and mRS [11], and DWIv and mRS [26]. The full (initial) model is shown in Figure 1.

**Figure 1:**
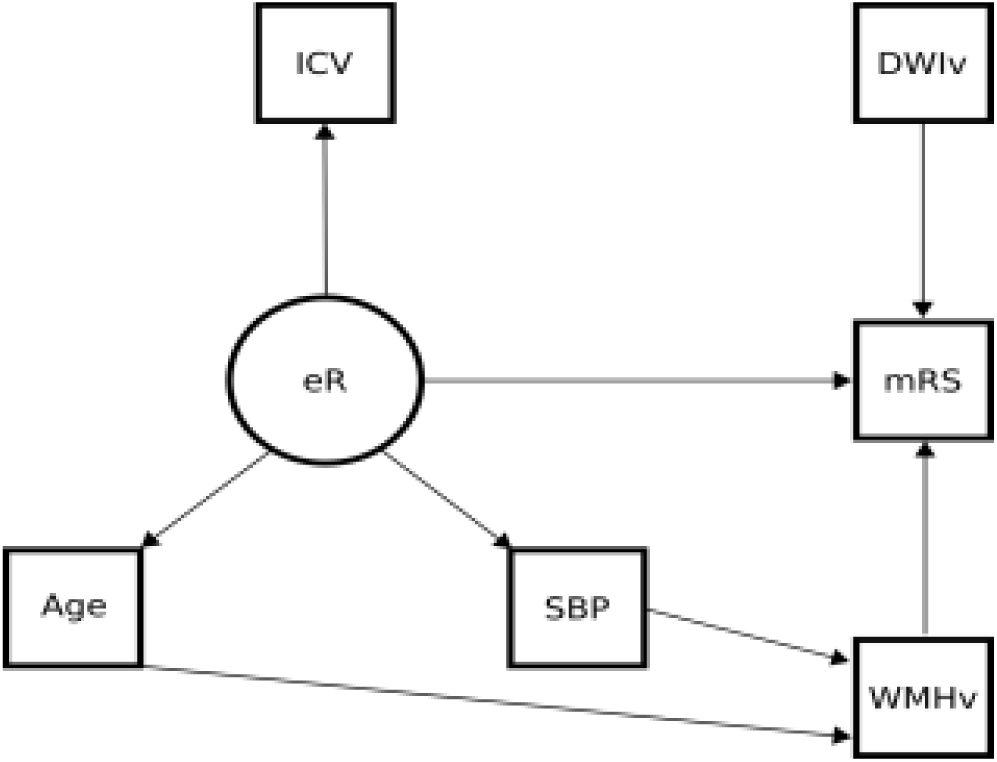
Initial structural equation model for effective reserve. *Squares represent measured phenotypes, whereas the circle indicates the latent variable. According to convention, arrows from latent variables point outwards. Associations in the model are indicated by arrows and can be estimated using path analysis.* Abbreviations: DWIv – acute lesion volume based on diffusion weighted images; eR – effective reserve; ICV – intracranial volume; mRS – modified Rankin Scale; SBP – systolic blood pressure; WMHv – white matter hyperintensity volume

### Model evaluation

Model parameters are estimated using the publicly available R package *LAVAAN* [27] for SEM estimation and path analysis without priors, using a maximum likelihood estimator with robust errors (MLR). The estimated path coefficients provide information on the magnitude and significance of the by the model hypothesized causal relationship between variables. After the estimation of the *full model*, non-significant factors/connections are eliminated resulting in a *simplified model*. It is common practice to use multiple statistical criteria to assess the model fit in SEM and to determine which model (full/simplified) best describes the data. In this work, these criteria (abbreviation; sign of a good model fit) are chi-square (χ^2^; low values, non-significant p-values), normed and comparative fit index (NFI/CFI; values greater than 0.95), root mean square error of approximation (RMSEA; values smaller than 0.5), Akaike information criterion for comparing models (AIC; smaller values indicate better fit) and Hoelter’s critical *n* (HCN; values greater than 200). Figure 1 describes the purpose of each of these measures. For a general overview and discussion of these metrics, see e.g. Kline[10].

Using the selected model, we first investigate the stability of the parameter estimation of the SEM model. In order to do so, we apply a 20-fold cross-validation, where the data are split in 20 approximately equally sized subsets and subsequently re-estimate the model based on 19 of those subsets. This is repeated 20 times and each time the estimated path-coefficients are recorded. Given the distribution for each parameter, we calculate the coefficient of variation as a summary measure.

Using the same 20-fold cross-validation approach, we estimate the parameters of the function given by equation (1) based on 19 folds. This allows us to estimate eR without inclusion of mRS on the remaining fold. Subsequently, we use the selected model and calculate the mRS score:

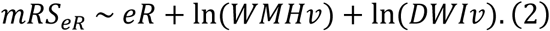

As a comparison, we use a linear model with the same folds where all observed variables (ICV, Age, SBP, WMHv and DWIv) directly relate to mRS (without introducing eR in the model):

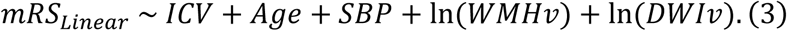

We then compare correlation coefficients between the observed mRS score and the estimates from equations (2) and (3) to evaluate the benefits of the added latent variable, i.e. eR.

### Analysis of hypertension status

Additionally, we use hypertension status (binary yes/no) in our data set as a proxy of the overall vascular health, where we assume better vascular health in those not exposed to long-term effects of hypertension (i.e., non-hypertensive patients), and investigate its relationship with eR. For this, we compare both groups with respect to the group-specific eR estimates produced by our model.

## Results

Baseline characteristics and stroke outcome of this cohort are summarized in Table 2. Here, non-hypertensive patients are on average older than hypertensive patients (72.0 years vs 64.4 years; p<0.001), while no difference in outcome is observed between these groups. Weak to moderate correlations are seen between most measures used in our model (Table 3).

**Table 1:** Assessment of model criteria for both full and the simplified model of post-stroke outcome.

| Criterion | Meaning | Full model | Simplified model |
| --- | --- | --- | --- |
| $\chi^2$ | Assesses magnitude of difference between the model and the data | 7.45 (p>0.38) | 7.59 (p>0.37) |
| NFI/CFI | Compares $\chi^2$ of the model to the “worst case scenario” (null hypothesis) | 0.96/1.00 | 0.96/1.00 |
| RMSEA | Estimates how well the model fits the population covariance matrix | 0.02 | 0.01 |
| AIC | Measure of relative quality of the models, given the data | 14530.14 | 14530.30 |
| HCN (alpha=0.05) | Indicative of adequate sample size | 856.93 | 840.99 |

Abbreviations: AIC – Akaike information criterion; HCN – Hoelter’s critical n; NFI/CFI – normed/comparative fit index; RMSEA – root mean square error of approximation

**Table 2:** Baseline characteristics and functional outcome of 453 acute ischemic stroke patients.

|  | All subjects | Hypertensive | Non-hypertensive |
| --- | --- | --- | --- |
| **Number of patients** | 453 | 311 | 70 |
| **Sex, n (%) male** | 165 (36.4) | 121 (39.0) | 24 (34.3) |
| **Age (years)\*\*\*** | 66.6 (55.7-77.6) | 64.4 (55.7-77.6) | 72.0 (47.7-73.6) |
| **ICV (cm<sup>3</sup>)\*** | 15.3 (14.5-16.1) | 15.3 (14.3-15.9) | 15.2 (14.4-17.2) |
| **Admission SBP (mmHg)\*\*\*** | 148.0 (132.0-169.0) | 148.0 (138.0-175.0) | 148.5 (126.5-157.8) |
| **ln(WMHv), (mm<sup>3</sup>)\*** | 1.7 (1.0-2.8) | 1.5 (1.1-2.8) | 1.9 (0.9-2.4) |
| **ln(DWIv), (mm<sup>3</sup>)** | 0.8 (-0.5-2.5) | 0.7 (-0.4-2.3) | 1.0 (-0.5-2.9) |
| **mRS [0-6]** | 1 (0-2) | 1 (0-3) | 1 (0-2) |

Each biomarker is reported as median (interquartile range). \*: p<0.05; \*\*: p<0.01; \*\*\*: p<0.001 for a test of differences between
hypertensive and non-hypertensive groups.
Abbreviations: DWIv – acute lesion volume based on diffusion weighted images; ICV – intracranial volume; ln – natural logarithm;
mRS – modified Rankin Scale; SBP – systolic blood pressure; WMHv – white matter hyperintensity volume

**Table 3:** Spearman correlation coefficients between clinical variables used in the model.

|  | ICV | ln(WMHv) | ln(DWIv) | SBP | mRS |
| --- | --- | --- | --- | --- | --- |
| **Age** | -0.24*** | 0.51*** | -0.02 | 0.22*** | 0.11* |
|  | **ICV** | -0.10* | -0.01 | -0.14** | 0.02 |
|  |  | **ln(WMHv)** | 0.02 | 0.20*** | 0.08 |
|  |  |  | **ln(DWIv)** | -0.05 | 0.11* |
|  |  |  |  | **SBP** | -0.02 |

\*, p<0.05; \*\*, p<0.01; \*\*\*, p<0.001
Abbreviations: DWIv – acute lesion volume based on diffusion weighted images; ICV – intracranial volume; ln – natural logarithm;
mRS – modified Rankin Scale; SBP – systolic blood pressure; WMHv – white matter hyperintensity volume

### Model estimation

Figure *2* shows the full and the simplified model, including their path coefficients with corresponding significance. Model parameters of the *full model* (left) imply that age and SBP negatively affect eR (path coefficients of -0.77 (p<0.01) and -0.29 (p<0.05), respectively), whereas ICV has a positive impact (p<0.01; path coefficient of 0.32).

**Figure 2:**
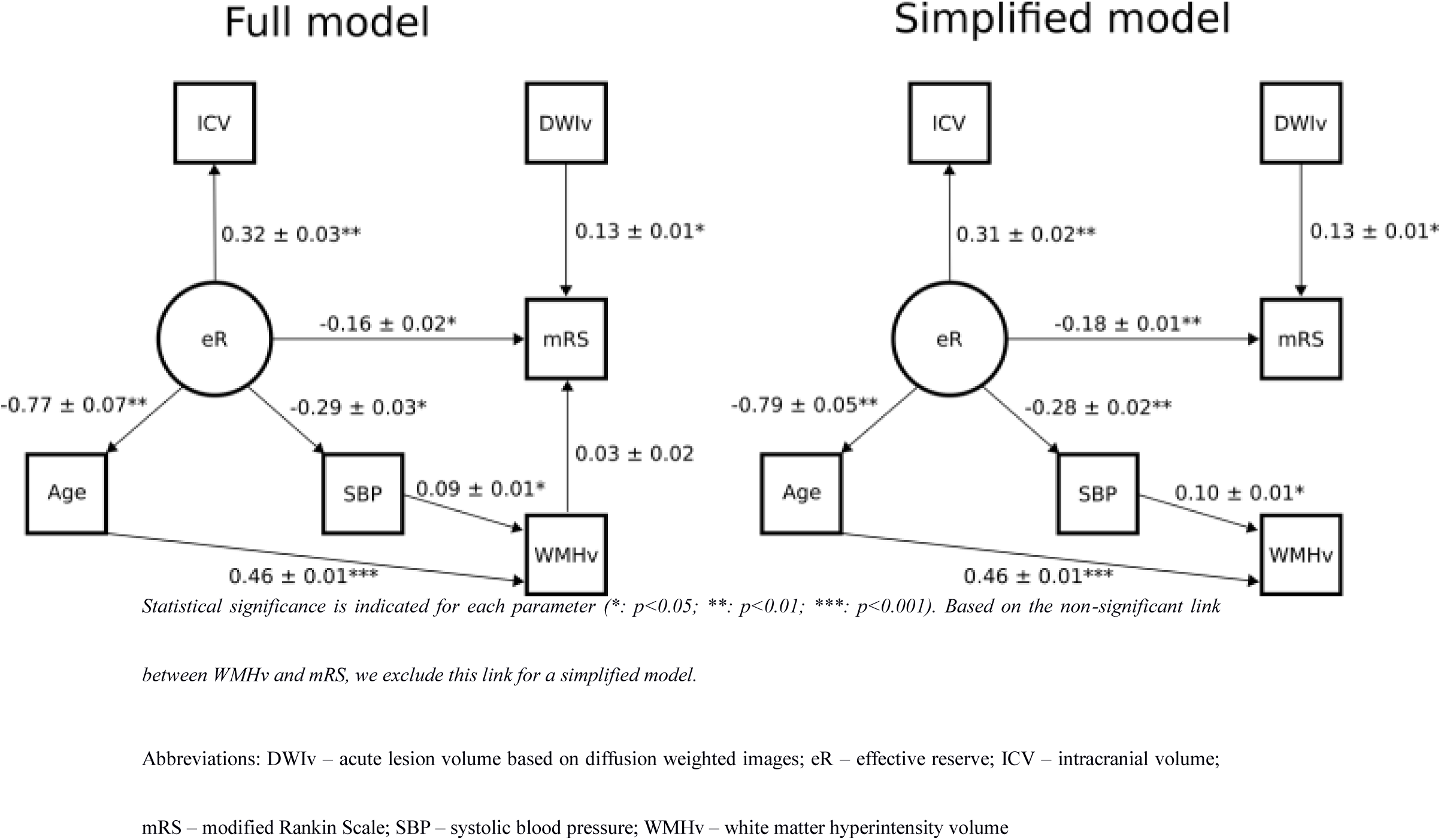
SEM with estimated association using path analysis.

The model exhibits significant positive association between SBP and WMHv (p<0.05; path coefficient of 0.09), and between age and WMHv (p<0.001; path coefficient of 0.46). DWIv was significantly associated with mRS (p<0.05; path coefficient of 0.13). Moreover, eR was negatively associated with mRS (p<0.05; path coefficient of -0.16), suggesting that a higher value of eR results in a lower value of mRS, reflecting a better outcome.

The full model suggests that there is no direct relationship between WMHv and mRS (p>0.05; path coefficient of 0.03). In a *simplified model* without this link (Figure *2*; right), path coefficients are nearly identical to those of the full model (on average the change is less than 5%) and statistical significance remains the same or improves (p<0.01 for both path coefficients between eR and mRS and between eR and SBP, compared to p<0.05 in the full model).

### Model selection

When assessed by the model evaluation criteria, the two models (full versus simplified) did not differ (Figure 1). Based on these results and the parsimony principle, we selected the simplified model for the remainder of our analysis.

Utilizing the 20-fold cross-validation, the absolute value of the coefficient of variation was on average below 8% (averaged over all parameters; see Figure *2*). Comparison of the observed mRS and each model’s estimate show that the calculated correlation coefficients were higher (mean ± SD; Fisher z-transformation p<0.001) for the model that incorporates eR (ρ = 0.29 ± 0.18; *mRS*_*eR*_) versus that of a linear baseline model (ρ = 0.15±0.17, *mRS*_*Linear*_)

### Analysis of hypertension status

There was statistical group difference in eR between non-hypertensive and hypertensive patients (Wilcoxon rank sum test p<10^−6^), where the non-hypertensive group exhibited higher eR.

## Discussion

BR has been proposed as a concept that corresponds to the brain’s capacity to withstand insults, and it has been shown to co-vary with WMHv [9]. In this analysis, we demonstrated that a latent construct, referred to as eR, may act as a surrogate measure of such a protective mechanism in AIS patients with regard to functional outcomes, i.e., higher eR is associated with a lower 90-day mRS. Moreover, the relatively low calculated coefficients of variation suggest that our model parameter estimations are stable, and that, based on correlations between estimated and observed mRS, the model which incorporated eR describes the data better than the baseline linear model. Furthermore, by splitting the stroke cohort into hypertensive and non-hypertensive patients, we demonstrated significant group differences with respect to their estimated measure of eR (higher eR in the non-hypertensive group). This supports the interpretation that eR may reflect a better ability to compensate for changes in vascular health, as approximated by hypertension status, or even a general protective mechanism.

Our results show that most input variables in our model are weakly to moderately correlated among each other. In spite of the correlation between WMHv and mRS in the data (ρ=0.1, p<0.05; Table 3), however, our full model did not indicate a significant direct association between the two. This is in contrast to other findings in the literature [11]. However, Brickman et al.[9] showed that their BR construct co-varied with WMHv while controlling for “outcome”, which means that it is more likely to see high WMHv in a patient with higher BR. In addition, our analysis suggests a slight, although insignificant, improvement when excluding the WMHv link to mRS. Therefore, it may be possible that the significant association between WMHv and mRS outcome score may be an observed effect of the association between eR and mRS instead. This might also suggest that eR mediates the previously observed correlations between WMHv and mRS or the other way around. Future studies are therefore warranted to disentangle this relationship.

We note that the path coefficient between age and eR, as shown in Figure *2*, was estimated with approximately twice the amplitude, compared to ICV and SBP. Similarly to the correlations with other observed variables (Table 3), age seems to be the strongest modifying factor for eR. The high negative impact may be representative of the amount of insult to human brains from which the brain has to recover over the lifespan. An investigation where patients are matched by age would be of interest and presents an interesting venue for future large scale studies.

Furthermore, we found a negative association between age and ICV (Table 3). This relationship may at first not seem intuitive for an adult cohort, as ICV has been shown to remain constant across age[28]. Moreover, the correlation persists after accounting for differences in sex (Spearman’s ρ -0.17, p<0.001). Considering that this is a cross-sectional study, one explanation could be a secular change in ICV [29]. Further studies are warranted to determine its origin.

There is a number of limitations to the presented study design. In this work, we used mRS, the most commonly used measure for stroke outcome in the literature [14]. However, it has been criticized for its inter-observer variability [30]. Moreover, mRS is not a continuous variable, which is one of the assumptions in the presented SEM model and most likely results in lower statistical significance in our analysis. This could further explain the observed non-significance between WMHv and mRS in our model. Adjusting the SEM software to accept ordinal variable classes could further improve future studies. Additionally, mRS summarizes differences in patients in predefined categories and is heavily weighted for motor function and ambulatory status. Using an outcome metric, which reflects more subtle functional differences, may further help improve the model and the investigation of eR as a protective mechanism in AIS.

The SEM approach models the shared variance between observations and can be used to estimate latent variables, for which no direct measurements exist, but its effects may be seen through other (indirect) measurements[10]. One of the strengths of SEM is grounded in its suitability for data with measurement errors or variability (e.g., inter-rater variations) or incomplete measurements (not all variables may be present in all subjects; [10]), which may be the case in clinical data. However, SEM is hypothesis-driven and is therefore not suitable for model generation, as each model relies heavily on experts to specify the links between the variables. Moreover, the more complex the model, the more data are required for it to converge and give an appropriate estimate of the relations within the model. The results in Table 1 (HCN) suggest that we have enough data to estimate each model parameter appropriately. However, the sample size limits the complexity and future directions, which can be taken using the current dataset. As an example, one bias in our results may be that we did not differentiate between male and female patients. It has been shown in the literature that many clinical variables are highly correlated with the sex of the patient [31], making this an interesting aspect of future work. However, in order to estimate the effects of sex on the estimated model parameter, which would require splitting the data and fitting an SEM model to each subset, a larger dataset is necessary.

In summary, we have introduced the concept of eR and have shown that it is inversely associated with functional post-stroke outcome, suggesting eR can be used as a descriptor of protective mechanisms in AIS patients. In particular, it may be representative of the vascular health and could therefore be of interest for future research and clinical applications.

## Acknowledgements

This study was supported by the NIH–National Institute of Neurological Disorders and Stroke (K23NS064052, R01NS082285, and R01NS086905), American Heart Association/Bugher Foundation Centers for Stroke Prevention Research, and Deane Institute for Integrative Study of Atrial Fibrillation and Stroke.

